# Conditional Knockout of Striatal Gnal Produces Dystonia-like Motor Phenotypes

**DOI:** 10.1101/2024.08.26.609754

**Authors:** Nicole E. Chambers, Dominic Hall, Stephanie Barsoum, Evan Miller, Tiffany Curry, Morgan Kaplan, Sarah Garan, Ignacio Gallardo, Rick Staab, Douglas Nabert, Kevin Hutchinson, Michael Millett, Mark S. Moehle

**Affiliations:** Department of Pharmacology and Therapeutics and Center for Translational Research in Neurodegeneration, University of Florida; Gainesville, Florida, 32610, United States of America

## Abstract

Loss-of-function mutations in *GNAL* have been linked to an adult-onset, isolated dystonia that is largely indistinguishable from idiopathic dystonia. *GNAL* encodes Gα_olf_, a heterotrimeric G-protein α subunit with a defined molecular function to increase the production of the second messenger cAMP. Gα_olf_ is abundant in the striatum, and is the only stimulatory G-protein in many cell types of the striatum. Due to the defined molecular signaling pathway and expression pattern of Gα_olf_, the clear genetic link to dystonia makes *GNAL* an exciting target to understand the pathological mechanisms of not only this genetic dystonia, but also the larger idiopathic disease. To better understand *GNAL*-linked dystonia, we generated a novel genetic mouse model that allows us to conditionally knock out *Gnal* in a site and time-specific manner. In the current study we used genetic or AAV based approaches to express Cre to knockout striatal *Gnal* in our novel *Gnal fl/fl* model. We then performed motor behavioral testing and *ex vivo* whole-cell patch clamp electrophysiology of striatal spiny projection neurons to interrogate how loss of *Gnal* leads to dystonia. Mice with conditional striatal knockout of *Gnal* show hindlimb clasping, other dystonia-like postures, less motor coordination, slowness, and torticollis as compared to age-matched controls. Furthermore, striatal spiny projection neurons show increased excitability in *Gnal* knockout animals. These exciting data are the first to report uninduced, overt dystonia in a mouse model of *GNAL-*linked dystonia, and directly correlate these with changes in spiny projection neuron electrophysiological properties. Our results show that adult loss of *Gnal* in the striatum leads to the development of dystonia, through homeostatic, paradoxical increases in spiny projection neuron excitability, and suggest that therapeutic strategies aimed at decreasing this hyperexcitable phenotype may provide symptomatic relief for patients with disease.

**One Sentence Summary:** When *Gnal* is knocked out in the striatum of mice we observe overt behavioral symptoms and hyperexcitability in striatal spiny projection neurons.

## INTRODUCTION

Dystonia is a group of hyperkinetic movement disorders which are associated with abnormal movements and muscle contractions(*1*). *GNAL*-linked dystonia (DYT25) is an adult-onset genetic form of dystonia which is caused by loss-of-function mutations in *GNAL*(*2–5*). *GNAL* encodes the α subunit of a heterotrimeric G-protein, Gα_olf_(*6–8*). Gα_olf_ is a stimulatory G-protein which causes the production of cAMP and activates the protein kinase A signaling cascade, and is similar in sequence and function to the classical stimulatory Gα protein Gα_S_(*6, 7, 9, 10*). Gα_olf_ is particularly enriched in the striatum, where Gα_olf_ is highly expressed in both D1 and D2 expressing striatal projection neurons (SPNs)(*11*). In SPNs, Gα_olf_ is the only stimulatory Gα subunit, and SPNs have low to no expression of Gα_S_ (*9, 11*). Other cell types in the striatum such as cholinergic interneurons have expression of both Gα_olf_ and Gα_S,_ or have no expression of Gα_olf_ (e.g. SST+ interneurons)(*11*). This suggests that Gα_olf_ may have a unique role in regulating SPN function. Within SPNs, Gα_olf_ couples to the dopamine D1 receptor in direct pathway SPNs (dSPNs) and the adenosine A2A receptor in indirect pathway SPNs (iSPNs)(*7, 8, 10*). Within the striatum, Gα_olf_ has been shown to be a critical regulator of SPN function in both normal physiological states as well as in other movement disorders(*5–8, 12–14*). However, how mutations in *GNAL* alter SPN function to cause dystonia remains unclear.

A major challenge in modeling *GNAL*-linked dystonia is that homozygous *Gnal* knockout in mice is postnatally lethal, and this is likely due to olfactory deficits in homozygous knockout mice(*6*). *Gnal* heterozygous mice are viable, but heterozygous mice do not show overt dystonia like postures(*6, 15–17*). However, dystonia like abnormal postures may be able to be induced through injection of the cholinergic agonist oxotremorine(*15, 17*). Using these mice and a similar rat model, multiple deficits in cholinergic interneuron firing, cortico-striatal plasticity, SPN firing, as well as cerebellar-thalamic-cortical network deficits have been shown(*2, 11, 15–22*). However, due to the lack of overt dystonia in these rodents, linking any of these deficits to the development of dystonia have been difficult. Furthermore, the global knockout model does not allow for testing the specific contribution of individual brain nuclei or neuron types to the development of dystonia.

Therefore, in the current study we created a novel model of *GNAL*-linked dystonia which inserted LoxP sites around exons 3 and 4 of *Gnal* to mimic loss-of-function mutations in *GNAL* which are causative for dystonia. Using either viral or embryonic driver line expression of Cre, we determined how loss of *Gnal* in the striatum leads to dystonia-like phenotypes. We investigated changes in motor behavior and striatal physiology using robust motor behavioral tests coupled with *ex vivo* electrophysiology of SPN activity. We show that our novel mouse model develops dystonia-like postures which correlate to changes in SPN activity. Our results suggest that *Gnal* loss produces homeostatic alterations in SPN activity and leads to the development of dystonia. Overall, the results of the current study may have implications on symptom management in patients with dystonia.

## RESULTS

### AAV-Conditional Knockout (AAV-cKO) Leads to Dystonia-Like Postures

To determine if loss of *Gnal* leads to dystonia, we injected *Gnal fl/fl* mice with an AAV expressing Cre or tdTomato as a control (Figure 1A). We confirmed that expression of the Cre virus led to loss of *Gnal* in virally infected striatal neurons (Figure S1A-C). AAV-cKO mice (control n = 28, AAV-cKO n = 25) suspended by their tail exhibit dystonic postures including forelimb clenching, hindlimb clenching, abnormal twisting of the trunk (Figure 1B), torticollis (Figure S2A), and hindlimb clasping (Figure 1C-1D, for guide to scoring hindlimb clasping see Figure S2B). AAV-cKO mice showed significantly more hindlimb clasping than control injected mice (Mann-Whitney U test, U = 14, p < 0.0001). Hindlimb clasping develops over weeks 5-8 after viral injection, (control n = 8; AAV cKO n = 7; Kruskal-Wallis test, ꭓ^2^ (8,92) = 79.18, p < 0.0001). AAV-cKO group showed more hindlimb clasping than controls for all time points listed (Dunn’s post-hoc test, Week 5, p = 0.04; Week 6, p < 0.001; Week 7-8, p < 0.0001), with hindlimb clasping reaching a plateau between 7-8 weeks after viral injection where hindlimb clasping is significantly more than on week 5 (Figure 1C, Friedman ANOVA, ꭓ^2^ (4) = 17.69, p = 0.0005, Dunn’s post-hocs, p = 0.018). This is consistent with the time that it takes for AAV-Cre to express and cleave exons 3 and 4 (Figure S3A-B). This hindlimb clasping phenotype is present in nearly all mice injected with AAV-cKO virus (Figure 1D). Once it develops, this hindlimb clasping phenotype is stable over time, with the same group of mice 6 months (control =8; AAV-cKO = 7) and 12 months (control = 8; AAV-cKO = 7) after injection showing similar amounts of hindlimb clasping (Figure S3C), and both groups showing more hindlimb clasping than controls (Kruskal-Wallis test, ꭓ^2^(6,52) = 44.16, p < 0.001, Dunn’s posthoc p< 0.01, and p < 0.001 for 6 months and 12 months, respectively). Additionally, the hindlimb clasping phenotype is present regardless of the age of mice at the time of injection (Figure S3D) with mice injected at 3-6 months (control n = 8; AAV-ckO n = 7) of age showing a similar phenotype to mice injected at 12-24 months of age ((control = 9; AAV-cKO = 13), ꭓ^2^(4,37) = 30.44, p < 0.0001). Hindlimb clasping is also not affected by sex of the mouse, with both male and female mice showing similar phenotypes (Figure S6E, control female n = 9; control male n = 18; AAV-cKO female n =12; AAV-cKO male n = 18), but both groups showing differences from controls, ꭓ^2^ (4, 57) = 36.99, p < 0.0001, Dunn’s post hoc (Control female vs AAV-cKO female, p = 0.0087, Control female vs AAV-cKO male, p = 0.0016, control male vs AAV cKO female and control M vs AAV-cKO M, p < 0.0001).

**Figure 1.**
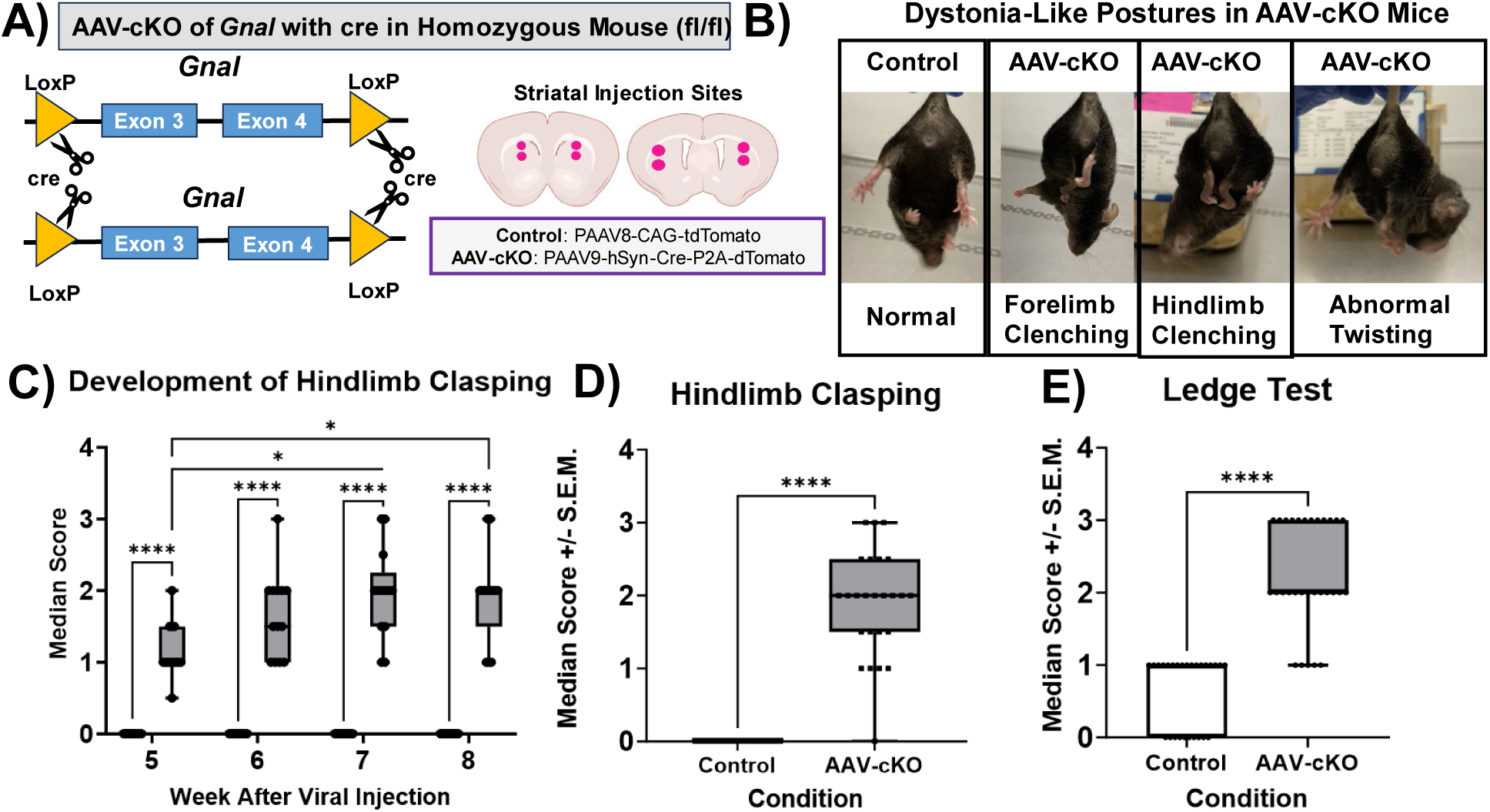
Viral *Gnal* Conditional Knockout (cKO) in the Striatum of *Gnal* floxed Mouse Leads to Dystonia-Like Postures. A) The mutation introduced in *Gnal* using CrispR-Cas is shown where LoxP sites flanking exons 3 and 4 were inserted. *Gnal* has germ line expression, thus there are two copies per mouse. LoxP sites allow for knockout of *Gnal* with a Cre virus (AAV-cKO). Consequences of cre-LoxP recombination are shown as scissors. Striatal slices from *BioRender* show injection sites of either control virus expressing just the fluorophore tdTomato, or the cKO virus containing cre and fluorophore dTomato. Viruses are listed for each condition. B) On the left is a normal control mouse with its hind limbs splayed out. On the right three panels, AAV-cKO mice demonstrate various dystonia-like postures including forelimb clenching, hindlimb clenching, and abnormal twisting. C) The timeline of development of a hindlimb clasping phenotype. These mice robustly develop dystonia-like symptoms by weeks 7-8 after injection. D) Shows median hindlimb clasping behavior in AAV-cKO mice. E) Shows median limb coordination on ledge test, lack of coordination is indicated by a higher score. Ledge test deficits are present in AAV-cKO mice. Data were analyzed using Mann Whitney U, α = 0.05.

We also correlated median hindlimb clasping scores with dystonic behaviors (Figure S4A-D; AAV-cKO, n = 10). Overall, we found that hindlimb clasping has no relationship with abnormal twisting behavior (Figure S4A, Spearman’s r = 0.13, p = 0.71). However, we found that hindlimb clasping is positively correlated with limb grabbing (Figure S4B, Spearman’s r = 0.90, p = 0.001), and forelimb clenching (Figure S4C, Spearman’s r = 0.71, p = 0.04). Furthermore, hindlimb clasping scores were positively correlated with time that the hindlimbs were together (Figure S4D, Spearman’s r = 0.70, p = 0.03). This suggests that hindlimb clasping is a valid proxy for dystonia-like movements.

To understand whether off target effects of the Cre-virus were influencing our interpretation of dystonia behavioral data, we performed a series of experiments to show optimal viral titers as well as any off-target cleavage effects of Cre. We used undiluted (1.0×10^13^ vg/ml) for AAV-cKO because diluted virus did not produce as robust of a phenotype in *Gnal* floxed animals (Figure S5A, Kruskal-Wallis test ꭓ^2^ (3,34) = 16.43, p < 0.001). To address whether this virus was producing a phenotype via *Gnal* knockout and not due to off-target effects, we injected wild-type C57BL6/J mice with either undiluted, 1.0×10^13^ vg/mL (n = 4), 3.33 x 10^12^ vg/mL (n = 5), or 1.0 x 10^12^ vg/mL (n = 5) of the AAV-cKO virus (pAAV9-hSyn,Cre-P2A-dTomato). We also injected *Gnal* floxed animals with AAV-control (pAAV8-CAG-tdTomato) in the same concentrations (1.0×10^13^ vg/mL (n = 25), 3.33 x 10^12^ vg/mL (n = 5), 1.0 x 10^12^ vg/mL (n = 4)) to compare with the wild type mice. Overall, there were no significant differences between groups of C57BL6/J mice and *Gnal* floxed controls, (Figure S5B, Kruskal-Wallis test, ꭓ^2^ (6,46) = 7.36, p = 0.20) indicating that the effects observed in the current study represent consequences of *Gnal* knockout and not random off-target effects.

### AAV-cKO leads to functional motor deficits

To determine if our AAV-cKO produced functional deficits resulting from dystonia-like motor behavior, we used the ledge test and Erasmus ladder(*23–27*). We observed that mice injected with AAV-cKO mice exhibit a lack of coordination in the ledge test as compared to AAV control mice, which likely represents a functional consequence of dystonia (control n = 27; AAV-cKO n = 30; Figure 1E, Mann-Whitney U, U(2) = 42.50, p < 0.0001, for ledge test scoring guide, see Figure S2C). Additionally, as with hindlimb clasping, ledge test scores do not change with the age of the mouse, (Figure S3E, ꭓ^2^ (4,57) = 37.16, p < 0.0001). There also are no sex differences in ledge test, although *Gnal* AAV-cKO mice of both sexes are different from controls (Figure S6F; female control, n = 11; male control, n = 17; female AAV-cKO, n = 10; male AAV-cKO, n = 15, Kruskal-Wallis test, ꭓ^2^ (4,53) = 43.20, p < 0.0001, female fl/fl vs Female AAV-cko = 0.0003, female fl/fl vs male gnal cKO, male fl/fl vs female Gnal cKO, and male fl/fl vs male cko < 0.0001). Development of the motor coordination phenotype mimics the timeline of development of hindlimb clasping, occurring 8 weeks after viral injection.

Using Erasmus ladder as a proxy of fine motor performance and gait (Figure 2A; control, n = 24; AAV-cKO, n = 18), we observed that AAV-cKO mice are slower in the ladder performance. Overall, AAV-cKO mice were slower to complete trials (Figure 2B, independent-samples t-test, t(40) = 2.4, p = 0.02). Mice were also slower to complete a variety of steps such as high rung backward steps, (Figure 2C, independent samples t-test, t(40) = 2.7, p = 0.01), high rung longsteps, (Figure 2D, independent samples t-test, t(40) = 2.9, p = 0.006), high to low rung longsteps, (Figure 2E, independent samples t-test, t(40) = 2.62, p = 0.01), high to low rung jumps (Figure 2F, independent samples t-test, t(39) = 2.69, P = 0.01), and high rung shortsteps, (Figure 2G, independent samples t-test, t(40) = 2.69, p = 0.01). Longsteps are steps in which mice cross more than one rung at a time, shortsteps are steps onto the next sequential rung. Overall, the results of this analysis suggests that mice have trouble moving normally due to functional consequences of dystonia-like behaviors.

**Figure 2.**
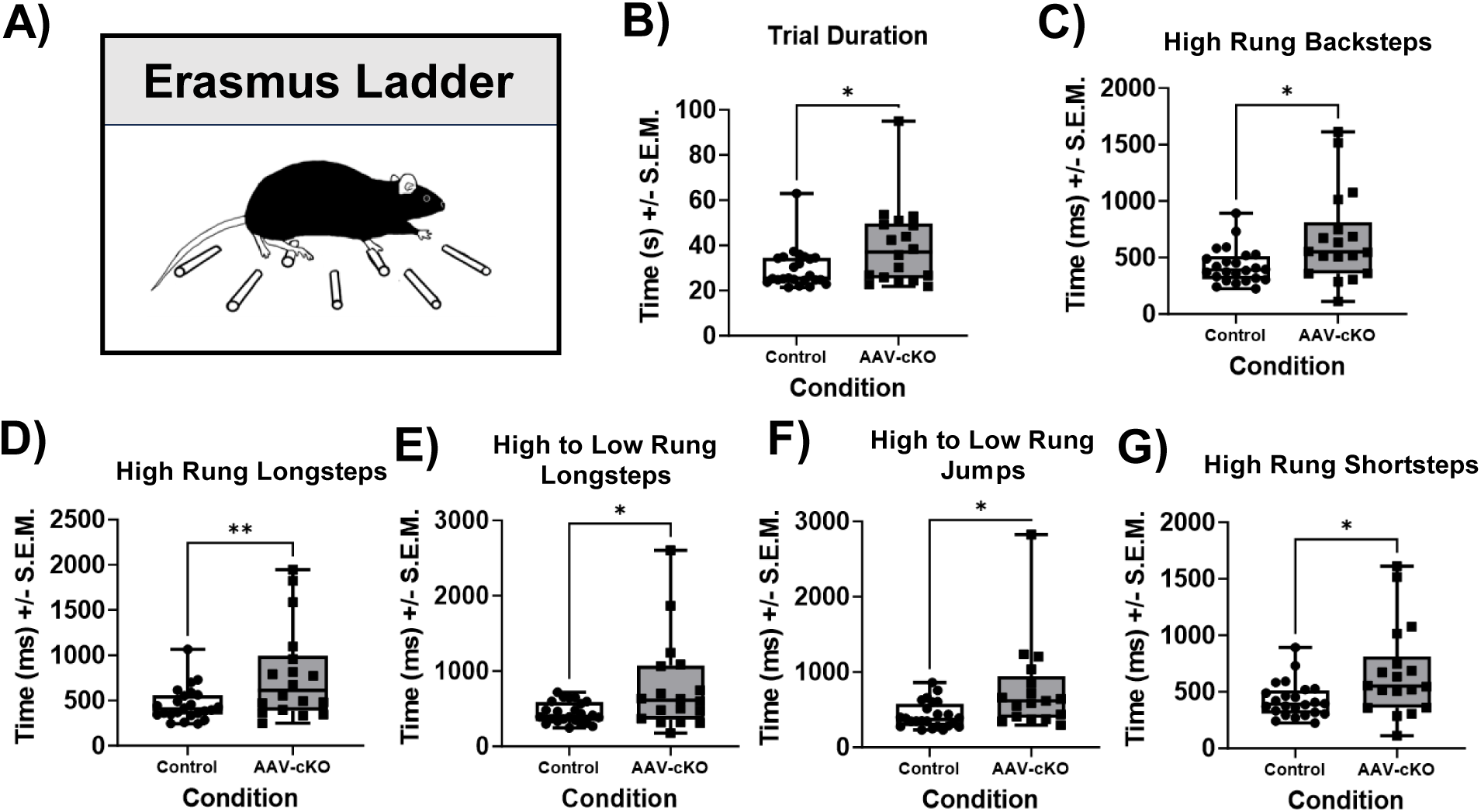
AAV-cKO Mice Show Slowness in Erasmus Ladder, Likely a Functional Consequence of Dystonia-Like Symptoms. A) A cartoon showing a mouse walking in the Erasmus ladder. There are high and low rungs for animals to traverse. Each of the data points shown on subsequent graphs represent an average was taken for one mouse for 42 trials walking across the ladder. All data were analyzed using two-tailed, independent samples t-test, α = 0.05. B) AAV-cKO mice take longer than control mice to complete a trial. C) AAV-cKO mice take longer than control mice to complete high rung backsteps within the ladder. D) Shows time to complete high rung longsteps with the ladder. AAV-cKO animals take longer than control mice to complete high rung longsteps. E) Shows time to complete high to low rung longsteps within the ladder. AAV-cKO mice take longer than control mice to complete high to low rung longsteps. F) AAV-cKO mice take longer than control mice to complete high to low rung jumps. G) Shows time to complete high rung shortsteps in the ladder. Overall, AAV-cKO mice take longer to complete high rung shortsteps than control mice.

Interestingly, we observed some sex differences in Erasmus ladder behavior, where female AAV-cKO mice do not show deficits in the Erasmus ladder in measures of fine motor performance and gait (female control, n = 6; male control, n = 18; female AAV-cKO, n = 5; male AAV-cKO, n = 13). For instance, female AAV-cKO mice completed trials in the Erasmus ladder in significantly less time than male AAV-cKO mice and were not different from controls (Figure S6A, one-way ANOVA F(3, 38) = 5.33, p = 0.0037, Tukey’s multiple comparisons p < 0.05). Similarly, female AAV-cKO mice completed high rung longsteps in less time than male AAV-cKO mice, and in similar amount of time as controls, (Figure S6B, One-way ANOVA, F(3,38) = 5.90, p = 0.0021, female fl/fl vs male AAV-cKO p = 0.0069, male fl/fl vs AAV-cKO p = 0.0084). Also, female AAV-cKO mice completed high to low rung longsteps in less time than male AAV-cKO mice, and in a similar amount of time to controls, (Figure S6C, One-way ANOVA, F(3,38) = 3.94, p = 0.0153 Tukey post hoc female fl/fl vs Male AAV-cKO p = 0.0377, male fl/fl vs Male AAV-cKO p = 0.0307). Finally, for high to low rung jumps, female AAV-cKO mice performed similarly to control mice and spent less time making high to low rung jumps than male AAV-ckO mice (Figure S6D, One-way ANOVA, female control, n = 6; male control, n = 18; female AAV-cKO, n = 5, male AAV-cKO, n = 12, F(3,37) = 3.35, p = 0.0292).Overall, these results suggest a possible protective effect of female sex on functional consequences of dystonia.

### Heterozygous *Gnal* floxed AAV-cKO mice show inconsistent hindlimb clasping and do not show motor coordination deficits in the ledge test

To determine how *Gnal* haploinsufficiency alters motor phenotypes in the AAV-cKO model, we injected heterozygous, *Gnal* fl/+ mice with AAV-cKO virus or control (Figure 3A-B). Overall, we found that AAV-cKO heterozygous, *Gnal* fl/+ mice show sparse hindlimb clasping. However, median hindlimb clasping scores between these mice and controls are not different (Figure 3C, Mann-Whitney U). Heterozygous *Gnal* fl/+ mice show more hindlimb clasping than controls when considering sums of hindlimb clasping behavior for 4 trials, suggesting that the incidence of hindlimb clasping is less in heterozygous, *Gnal* fl/+ AAV-cKO mice than in homozygous *Gnal* fl/+ mice (Figure3D, Mann-Whitney U, U = 0, p = 0.008).

**Figure 3.**
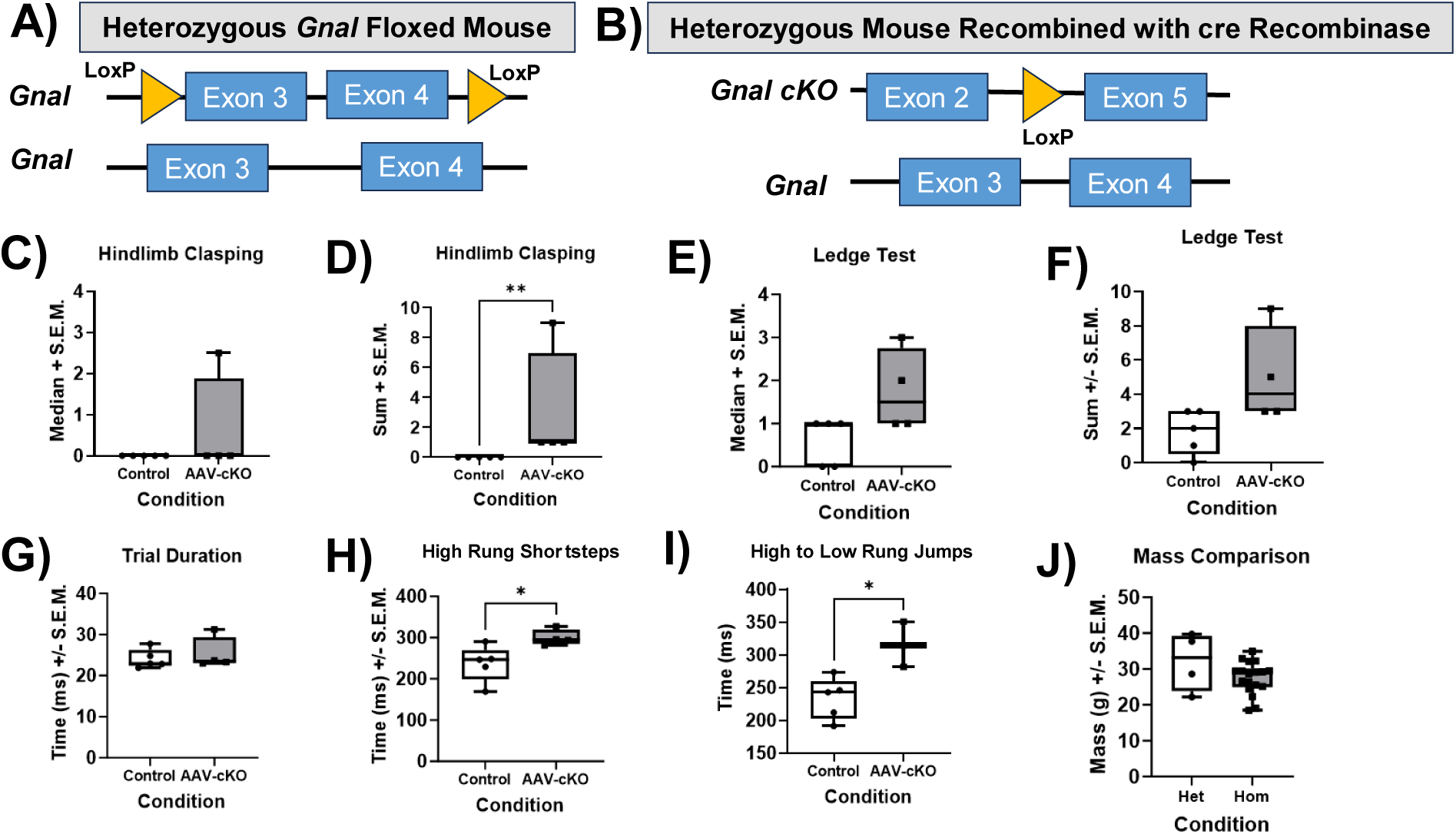
Heterozygous, *Gnal* fl/+ Mice show inconsistent hindlimb clasping and functional motor deficits in the Erasmus Ladder. A) Shows the one floxed copy of *Gnal* in heterozygous mice. B) Shows the consequences of knocking out *Gnal* with a Cre virus. C) Heterozygous AAV-cKO mice show no difference in median hindlimb clasping. D) Hetereozygous AAV-cKO mice show more hindlimb clasping than controls but only for sums of 4 trials. E) heterozygous AAV-cKO mice show no difference in motor coordination on ledge test than for controls whether analyzing medians as in E) or sums as in F). G) Trial duration is unchanged from that of controls in the Erasmus ladder. H) heterozygous AAV-cKO mice show longer time to complete high rung shortsteps than controls. I) Heterozygous AAV-cKO mice show longer time to complete high to low rung jumps than controls. J) Mass is not different between heterozygous AAV-cKO mice and controls. All data in this figure were analyzed using two-tailed, independent samples t-tests to evaluate differences between mice (α = 0.05).

### Heterozygous *Gnal* floxed AAV-cKO mice show functional deficits in Erasmus ladder but not in ledge test

To determine how *Gnal* haploinsufficiency alters functional motor deficits in the AAV-cKO model, we injected heterozygous, *Gnal* fl/+ mice with AAV-cKO virus or AAV-control virus and tested them on the ledge test and Erasmus ladder. Heterozygous, *Gnal* fl/+ mice (control, n = 5; AAV-cKO, n = 4) show reduced coordination in the ledge test. However, neither median scores of ledge test or sums for heterozygous AAV-cKO mice were not different controls (Figures 3E-F, Mann-Whitney U, non-significant). This suggests that motor coordination deficits are not present in heterozygous *Gnal* fl/+ mice.

Similarly, in the Erasmus ladder, we found that AAV-cKO heterozygous, *Gnal* fl/+ mice show no difference in overall trial duration (Figure 3G, two-tailed, independent samples t-test, non-significant) showing that overall functional deficits are not severe enough to interfere with global fine motor control. However, *Gnal* fl/+ AAV-cKO mice were significantly slower than controls on High rung shortsteps (Figure 3H, t(7) = 2.62, p = 0.034). Additionally *Gnal* fl/+ AAV-cKO mice were significantly slower than controls in the time that it took them to complete high to low rung jumps (Figure 3I, two-tailed, independent-samples t-test, t(6) = 3.44, p = 0.014).

Finally, we compared the mass of heterozygous AAV-cKO mice with that of heterozygous controls and found no differences in body mass, (Figure 3J).

### *Gnal* embryonic knockout (e-cKO) with *Rgs9* cre mice produces similar dystonic postures and hindlimb clasping to AAV-cKO

To evaluate the effects of knocking out *Gnal* embryonically in heterozygous *Gnal* floxed mice (Figure 4A, Figure S7B) or knocking out *Gnal* embryonically using homozygous *Gnal* floxed mice (Figure 4B, Figure S7A), we crossed homozygous *Gnal* floxed male mice with heterozygous *Gnal floxed Rgs9* cre female mice to produce an embryonic conditional knockout (e-cKO)(*28*). Similar to AAV-cKO mice, homozygous e-cKO mice show hindlimb clasping whereas control mice do not (Figure 4C, Mann-Whitney U, U(4) = 16.87, p = 0.0008). Interestingly, unlike AAV-cKO mice, heterozygous e-cKO mice do not show hindlimb clasping which is significantly greater than that of control mice, suggesting that embryonic mice are able to compensate for heterozygous e-cKO. Our e-cKO mice also show dystonia-like postures which mirror those of AAV-cKO mice, with front limb clenching, hindlimb clenching, twisting, and hindlimb clasping (Figure 4F). When comparing hindlimb clasping scores of heterozygous and homozygous *Gnal* e-cKO to homozygous AAV-cKO, there is no statistical difference between homozygous *Gnal* e-cKO and homozygous AAV-cKO; however, heterozygous e-cKO mice show less hindlimb clasping than AAV-ckO mice (Figure S8A, Kruskal-Wallis, ꭓ^2^ (3) = 12.40, p = 0.002).

**Figure 4.**
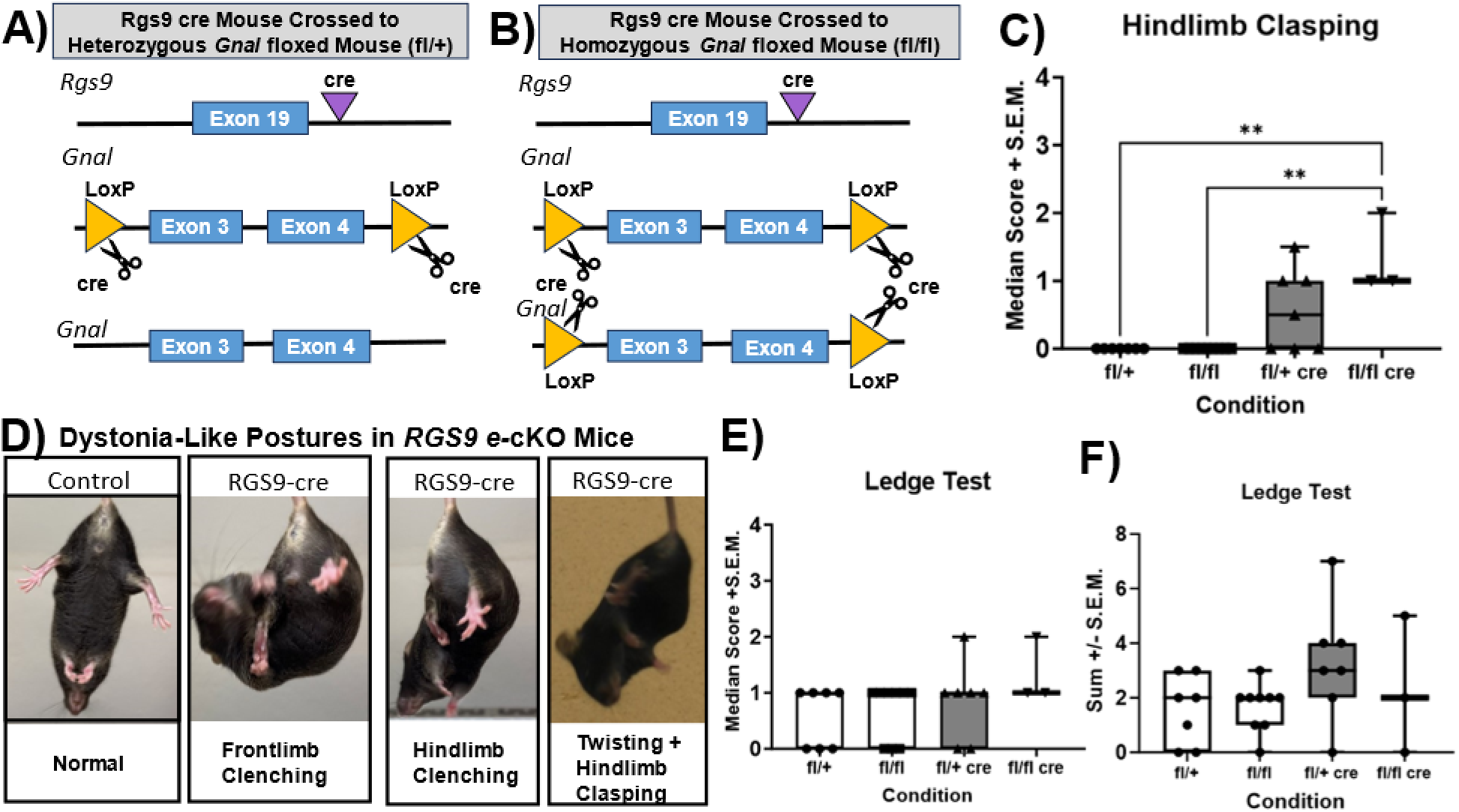
Embryonic Striatal Knockout (e-cKO) Produces a Similar Dystonic Phenotype to AAV-cKO. A) Shows genetic modifications in mouse heterozygous for *Gnal* flox gene (fl/+) that expresses *Rgs9* Cre. *Rgs9* is a regulator of G-protein signaling gene which is mainly located in the striatum, providing relatively selective *Gnal* knockout in Rgs9-cre animals. B) Genetic modifications in mouse homozygous for *Gnal* flox gene (fl/fl) that expresses *Rgs9*cre; both copies of *Gnal* are modified and subsequently deleted. C) Shows median scores for hindlimb clasping for mice heterozygous and homozygous for *Gnal* flox gene and negative for Cre (controls) plus heterozygous and homozygous mice which are positive for *Rgs9* Cre gene. Both heterozygous and homozygous controls display less hindlimb clasping than homozygous Cre + animals. D) Shows median scores for mice for ledge test, there are no significant differences between groups. E) Similar to AAV-cKO mice, Rgs9 Cre e-cKO mice show abnormal dystonia-like postures when suspended by their tail. The first image shows a normal, control mouse with limbs splayed out. Mice positive for Rgs9 Cre show abnormal front limb and hind limb clenching, as well as twisting and hindlimb clasping. For all data analysis Kruskal-Wallis ANOVA was used (α = 0.05).

### *Gnal* e-cKO mice do not show deficits in ledge test but do show functional deficits in the Erasmus Ladder

To measure functional deficits caused by dystonia-like behaviors in the e-cKO heterozygous and homozygous mice we used the ledge test and Erasmus ladder. Unlike AAV-cKO mice, neither homozygous nor heterozygous mice show a significant difference in coordination on ledge test as compared to control mice, and this is true whether comparing medians (Figure 4E, Kruskal-Wallis, non-significant), or sums, (Figure 4F, Kruskal-Wallis, non-significant). Furthermore, when comparing median scores of ledge test, homozygous AAV-cKO shows less coordination in ledge test than heterozygous e-cKO, with no difference between AAV-cKO and homozygous e-cKO (Figure S8B, ꭓ^2^ (3) = 10.75, p = 0.005).

To measure functional fine motor and gait deficits we used the Erasmus ladder. Overall, *Gnal* e-cKO mice had an increased trial duration as compared to control mice, showing that like AAV-cKO mice, they have functional deficits resulting from *Gnal* knockout (Figure S7C, independent-samples t-test, t(11) = 2.25, p < 0.046). Additionally, *Gnal* e-cKO mice show longer time to complete a variety of steps including low to high rung longsteps, (Figure S7D, independent samples t-test, t(11) = 2.22, p = 0.0483), and high to low shortsteps with their left foot, (Figure S7E, independent samples t-test, t(11) = 2.23, p = 0.0475). Finally e-cKO mice showed less high rung longsteps with their left forepaw, t (11) = 2.63, p = 0.02, indicating that these mice were likely taking more short steps with their forepaw, indicating that they may have been taking smaller steps overall, (Figure S7F, independent samples t-test, t(11) = 2.25, p = 0.04).

### *Gnal* e-cKO mice are not a viable model due to postnatal lethality and low body weight

Despite breeding over 60 litters of mice (*Rgs9* cre positive *Gnal* floxed heterozygous females bred to homozygous *Gnal* floxed males), we were only able to obtain and test 10 e-cKO animals which were included in this manuscript (heterozygous n = 7, homozygous n = 3). Although mice were born in normal numbers, *Gnal* e-cKO appears to be postnatally lethal. This is similar to past research which shows that global embryonic knockout of *Gnal* is postnatally lethal, likely due to olfactory dysfunction. In the current study we observed that mouse weights were particularly low for *Rgs9* Cre positive females on the *Gnal* fl background (Figure S8C). This suggests that animals positive for *Rgs9* cre and *Gnal* flox were less motivated to feed. Additionally, mice born to *Rgs9* cre positive dams on the *Gnal* floxed background showed overall lower body weight than homozygous *Gnal* floxed controls, (male controls born to Rgs9 cre mothers vs. *Gnal* floxed male controls p = 0.01). Taken together, our results indicate that embryonic knockout with *Gnal* is not a viable strategy for knocking out *Gnal* due to postnatal lethality and low body mass in surviving e-cKO mice.

### Loss of *Gnal* alters intrinsic properties of striatal SPNs

To understand physiological changes which occur in AAV-cKO mice, we performed surgery in which striatal injections were performed on the left side of the brain with the control AAV, and the right side of the brain with the AAV-cKO virus (Figure S9A). To examine effects of *Gnal* cKO and to leave G-protein signaling intact, we performed perforated patch recordings with amphotericin B (for procedure, see Figure S9D). Spiny projection neurons were identified by their electrophysiological characteristics including resting membrane potential, responsivity to current injection, and lack of spontaneous firing, and we only patched onto neurons positive for AAV-cKO or control-linked fluorophore. Following membrane perforation by amphotericin B (verified in membrane test) we recorded the following measures of intrinsic properties: membrane capacitance, membrane resistance, tau, resting membrane potential, and rheobase current. Using independent samples t-tests, we compared the membrane properties of SPNs of homozygous AAV-cKO mice with those of homozygous AAV-control mice to assess the effects of Gnal knockout on cell physiology. There was no change in cell capacitance, membrane resistance, tau, or resting membrane potential (Figure 5A-D). However, AAV-cKO decreased rheobase current, suggesting that cells in which *Gnal* is knocked out are hyperexcitable (Figure 5E, independent-samples t-test, t(15) = 2.66, p = 0.018).

**Figure 5.**
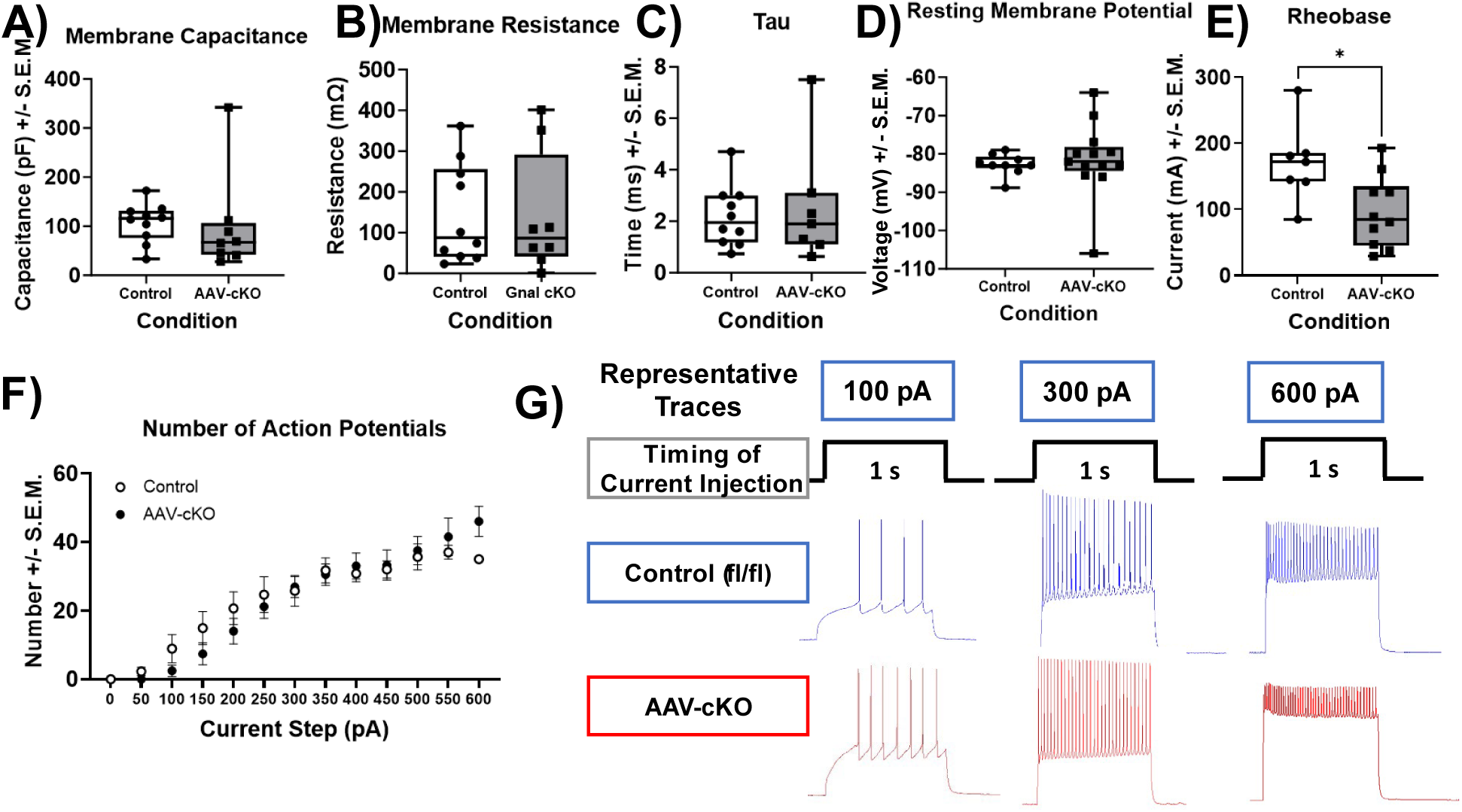
AAV-cKO Changes Lowers Rheobase, Does Not Affect Other Intrinsic Properties or Response to Current Injection in Spiny Projection Neurons. For each displayed graph in figure 5A-E two-tailed, independent samples t-tests were used to evaluate differences between groups, α = 0.05. Each dot represents a cell. A) Shows membrane capacitance (in picofaraday, pF) for control spiny projection neurons (SPNs) was not different for control and AAV-cKO cells. B) Membrane resistance (in milliOhms, mΩ) was not different for control and AAV-cKO cells. C) Tau (ms) for control and AAV-cKO SPNs was not different. D) Resting membrane potential (in millivolts, mV) was not different between control and AAV-cKO SPNs. E) Rheobase (amount of current to generate an action potential, measured in milliAmps, mA) was significantly lower in AAV-cKO than in control SPNs, suggesting that AAV-cKO SPNs are hyperexcitable. F) Number of evoked action potentials during current clamp with 1s of applied current (−350 to 600, in steps of 50 pA), analyzed by mixed-effects model shows no difference in the overall number of action potentials between control and AAV-cKO SPNs. G) Representative traces of SPN activity

### Loss of *Gnal* does not affect overall response to current injection

Next, to determine the extent to which SPNs are hyperexcitable, we performed input-output curves of depolarizing current steps and measured elicited action potentials. To do so we used 1s current injections which occurred every 20 s starting at -350 pA and increasing to 600 pA by steps of 50 pA. Overall number of action potentials were unchanged across conditions but the number of action potentials was greater for cells in both conditions across increasing current steps (Figure 5F, mixed effects analysis, non-significant). Representative traces for control and AAV-cKO are shown in Figure 5G. Peak-to-peak time and peak-to-peak frequency also remained unchanged (Figure S8B-C, mixed effects analysis, non-significant).

### AAV-cKO mice show changes in characteristics of action potential

In order to further examine differences in electrophysiological properties exhibited by loss of *Gnal*, we examined waveform characteristics of action potentials. For grand average action potentials see Figure 6A, for phase plots see Figure 6B. Overall, AAV-cKO animals show altered waveform characteristics compared to controls. We found that tau rise was increased in AAV-cKO animals, (Figure 6C, independent samples t-test, t(19) = 3.11, p = 0.005). Notably, tau rise represents the activity of sodium channels, suggesting that there is a difference in channel kinetics that results from the knockout. We did not see differences in tau(decay), Vmax, Vthreshold, or half-width (Figures 6D-G).

**Figure 6.**
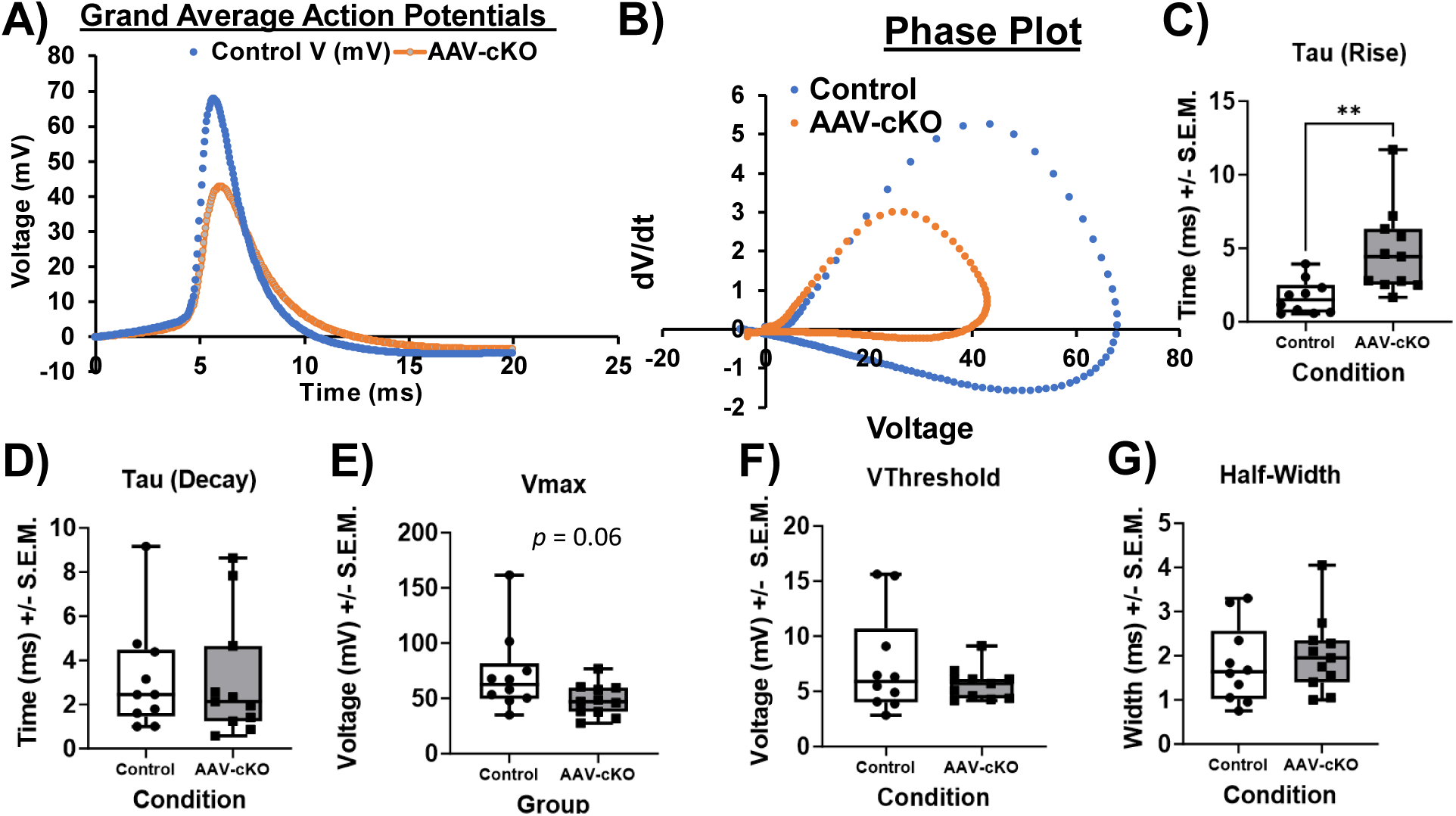
AAV-cKO Mice Show Differences in Characteristics of Action Potential. A) Shows grand average of action potentials evoked during all current steps before depolarization block for both control (blue) and AAV-cKO (orange). B) Shows the grand average phase plot of the action potentials for control and AAV-cKO. Overall, the phase plot shows that there are likely differences in multiple properties of the action potential. C) Tau rise (ms) was increased in AAV-cKO mice relative to controls. D) Tau decay was not different between control and AAV-cKO. D) Shows the maximum voltage (Vmax in millivolts, mV), while this approached significance (p = 0.06), there was no significant difference between control and AAV-cKO groups. E) Threshold voltage (Vthreshold, measured in mV) was not signicantly changed by AAV-cKO. G) Action potential half-width (ms) was not significantly changed in AAV-cKO mice vs. controls. All data were analyzed using two-tailed, independent-samples t-tests (α = 0.05). Each dot graphs C-G represents one cell.

### Dystonic symptoms in AAV-cKO mice are correlated with electrophysiological properties

In order to examine how changes in electrophysiological properties in SPNs related to dystonia-like behavior, we performed correlation analyses on these data. We observed a positive correlation between hindlimb clasping behavior and rheobase, (Figure S10A, Spearman’s r = 0.78, p = 0.03). Similarly, we observed a negative correlation with hindlimb clasping score and tau, (Figure S10B, Spearman’s r = -0.88, p = 0.03). Together this suggests that alterations in electrophysiological properties may drive dystonia-like behaviors.

## DISCUSSION

It has historically been challenging to investigate how loss-of-function mutations in *GNAL* result in dystonia symptoms. This is due to postnatal lethality in global *Gnal* knockout rodents(*6*), or the need to inject cholinergic agonists to elicit dystonia-like symptoms in existing models(*15, 17*). These challenges have limited the translational impact of existing models. Our new model showing the development of overt dystonia in response to viral-mediated, adult knockout of *Gnal* in the striatum may represent an exciting tool to understand the pathophysiology of *GNAL-*linked dystonia and the larger idiopathic disease. Our model exhibits dystonia-like postures that are strikingly similar to those of humans with *GNAL-*linked dystonia(*2–5, 29–35*). Specifically, we observed limb postures and torticollis in our mouse model reminiscent of human disease. Additionally, similar to human *GNAL*-linked dystonia, once symptoms have developed in our model they remain stable over time. Together, this mouse represents an advancement in our ability to model this genetically-linked dystonia.

Here, we demonstrated both functional behavioral consequences of striatal *Gnal* knockout and correlated electrophysiological changes which resulted from adult AAV-cKO in SPNs. This suggests that striatal dysfunction may be an important driving factor in the development of dystonia. However, we have recently shown that Gα_olf_ is expressed throughout many neuron types and brain nuclei, and is most abundantly expressed in Purkinje Cells of the cerebellum. This raises the possibility that other brain nuclei and neuron types are also capable of producing dystonia, and future studies to determine if the striatum is sufficient or necessary for the development of dystonia will be required.

Our data also demonstrate that adult loss of *Gnal* activity in the mouse appears to be critical to the development of robust dystonia-like phenotypes. While both adult heterozygous loss of *Gnal* and embryonic heterozygous or homozygous loss of *Gnal* in the striatum driven by *RGS9*-Cre led to abnormal movements, only adult homozygous AAV-cKO led to a robust dystonia-like phenotype. Adult homozygous loss of *Gnal* may be necessary for the development of dystonia, as the adult knockout allows for the escape of post-natal lethality and failure to thrive phenotypes observed with homozygous embryonic loss; while allowing for a sufficient homeostatic alteration of SPN activity and downstream behavior that is not driven by only partial adult *Gnal* loss.

We make several interesting novel observations with our striatal *Gnal* model. First, we show that hindlimb clasping, often used as a proxy measure of dystonia in mice, correlates with the expression of overt dystonic postures in this model, except trunk twisting. This suggests that hindlimb clasping is a useful proxy of dystonia symptoms, and that trunk twisting may occur through separate mechanisms or pathways to other dystonia-like postures. Second, we show that although male and female mice exhibit overt dystonia-like behaviors, they show differential functional motor consequences. Male mice were severely more impaired in Erasmus Ladder measures of fine motor performance than female mice showing similar levels of dystonic postures. How these sex differences arise in functional consequences necessitates further investigation.

Interestingly, we see a paradoxical increase in excitability in SPNs that lack *Gnal*. How the loss of a stimulatory G-protein subunit leads to a paradoxical increase in excitability, and not a decrease in excitability as we predicted, is not understood. This may suggest that SPNs undergo homeostatic alterations in intracellular signaling or channel kinetics as suggested by our alterations in action potential characteristics, especially Tau(rise). Additionally, it may be possible that the Gβγ subunits that are normally sequestered by Gα_olf_ may now be freely active, and contribute the altered excitability of SPNs(*20*). Either of these outcomes leads to implications for novel treatments for patients with *GNAL* linked dystonia. Our electrophysiological alterations correlate with the extent of hindlimb clasping and Tau(intrinsic), suggesting that the extent to which SPNs are affected drives symptoms. Normalizing this excitability back to baseline may therefore mitigate dystonia symptoms, and would implicate novel therapeutic strategies for the treatment of disease. Use of selective cholinergic agonists, possibly of M_4_, rather than broad spectrum cholinergic antagonists currently used clinicallly may ameliorate disease symptoms(*36–39*). This would be similar to other hyperkinetic movement disorders such as dyskinesia(*40*).

Throughout this study, we utilize AAVs that cannot discriminate between striatal neuron types. We have previously shown that most striatal neuron types express Gα_olf_, except for SST+ striatal interneurons(*11*). While SPNs are >90% of all neurons in the striatum, we cannot rule out a microcircuit-based effect driven by cholinergic interneurons or parvalbumin interneurons being a key driver of disease(*41*). As shown by our *RGS9* experiments, using Cre driver lines does not recapitulate disease well as confounding factors with feeding and other non-dystonia linked phenotypes influence interpretation of these experiments, and necessitate different novel genetic approaches. Development of FlpO based lines, to then drive the expression of Cre in specific striatal neuron subtypes, will be key to fully understand and dissect the role of individual neuron types in the development of dystonia.

This is the first model of *GNAL*-linked dystonia to show an overt behavioral phenotype which has face validity with the human condition. For the first time, this model allows us to examine the functional consequences of *Gnal* knockout in specific brain structures and in specific neuronal subtypes, and has allowed for the direct linkage in changes in electrophysiological properties with dystonia-like behaviors. Overall, this new model provides exciting opportunities for translational discoveries and rational design of novel therapeutic strategies.

## MATERIALS AND METHODS

### Study Design

The objective of the current study was to examine a new model of *Gnal*-linked dystonia which was created using a novel transgenic mouse—the *Gnal* floxed mouse line with LoxP sites flanking exons 3 and 4 (Figure 1A). To generate controls and conditional *Gnal* knockout mice, we first injected AAV-control virus (AAV8-CAG-tdTomato) or AAV-cKO virus (pAAV9-hSyn-Cre-P2A-tdTomato). We then characterized dystonia-like behaviors and functional consequences of dystonia in these animals over time. Next we investigated embryonic knockout (e-cKO) using mice which were either homozygous (fl/fl) or heterozygous (fl/+) for *Gnal* flox crossed with Rgs9 cre mice. Overall, we observed similar behavioral consequences in e-cKO animals as in our AAV-cKO. We also studied the effects of heterozygous *Gnal* floxed mice which were injected with an AAV-control or AAV-cKO virus, creating a *Gnal* conditional Knockout (AAV-cKO). These animals also showed similar deficits, but symptoms were less frequent than in AAV-cKO mice. Finally, we characterized physiological activity in striatal spiny projection neurons (SPNs) in response to AAV-cKO. To do so, we injected one half of the striatum with the AAV-cKO virus, and the other half of the striatum with the AAV-control virus. We euthanized the animals, prepared the tissue, and we performed perforated patch electrophysiology recordings with amphotericin B in K-Gluconate internal. Perforated patch recordings were used because *Gnal* encodes the alpha subunit of the heterotrimeric G-protein Gα_olf_, and using perforated patch allows us to keep this G-protein and relevant internal cell properties intact. This is because in perforated patch we only seal onto the cell membrane and allow time for small perforations to form which allows for electrical access to striatal SPNs without washing out the contents of the cell. SPNs were identified by their size and electrophysiological properties (resting membrane potential -80 to -85 mv and no spontaneous activity). We examined both intrinsic properties of the neuron (membrane capacitance, membrane resistance, tau, resting membrane potential, and rheobase) as well as response to current injection (−350 pA to 600 pA, 50 pA steps). We then used the action potentials generated through current steps to investigate differences in properties of the action potential.

**Subjects:** For experiments involving homozygous *Gnal*-floxed mice, *Gnal*-floxed mice on a C57BL/6J background underwent surgery beginning at 3 months of age. In one of the experiments mice underwent surgery at 12 – 24 months of age (see S3). For experiments involving heterozygous *Gnal* floxed mice, homozygous *Gnal* floxed mice were bred to cre-driver lines and resulting offspring that did not express cre were used. Notably, only female cre mice were bred to male *Gnal* floxed mice to prevent off-target effects of cre.

For experiments involving *Rgs9* cre mice(*28*), female *Rgs9* cre mice were bred to *Gnal* floxed male mice, and resulting offspring were earpunched, genotyped, and used in experiments. Mice used in *Rgs9* cre experiments did not undergo surgery as cre was induced embryonically via genetic expression.

All mice were kept in a colony room maintained at 24 degrees on a 12 h light-dark cycle (lights on at 0700). Mice were provided with ad libitum access to food and water and were kept in group housing in groups of up to 5 mice in plastic cages. All procedures were performed in accordance with the IACUC at the University of Florida and with the current version of *The Guide for the Care and Use of Laboratory Animals.* **Surgery:** Mice were placed in an anesthesia chamber and anesthetized with 2-3% isoflurane mixed in oxygen. Mice were weighed, their scalp shaved, and then they were injected with a pre-operative dose of carprofen (5 mg/kg, s.c.) and ropivacaine (300 microliters) was injected into the scalp. Mice were stabilized in an RWD stereotax and then maintained at 1-2% isoflurane in oxygen throughout the surgery. Alternating scrubs of chlorhexidine and saline were used to clean the surgical site. Next a scalpel was used to cut through the scalp and expose bregma. Target coordinates were identified according to bregma. Next, a burr hole was drilled carefully over each of 4 separate injection sites. A total of 8 striatal injections (0.5 μL) were performed at the following coordinates: (Anterior coordinates: AP: 1.20 mm; ML: ±1.25 mm; DV: -3.50 mm and - 2.50 mm; Posterior coordinates: AP: 0.40 mm, ML: ± 2.40, DV: -3.50 mm and -2.50 mm) and mice received either control AAV9-hSyn-tdTomato or dystonia virus AAV9-hSyn-cre-p2A-tdTomato. Following surgery mice received an injection of saline to stave off dehydration. Furthermore, mice were given moistened chow and gel cups as well as postoperative carprofen every 24 h for 48 h following surgery. For the electrophysiology surgeries, the right side only was injected with AAV9-hSyn-cre-p2A-tdTomato and the left side was injected with control virus, AAV9-hSyn-tdTomato.

### Breeding Scheme

Unless otherwise stated mice were bred from the combination of Gnal floxed parents. For the embryonic knockout RGS9 cre female mice were bred with Gnal floxed males. Heterozygous fl/+ animals were generated from the combination of *Rgs9* cre females crossed with male *Gnal* floxed mice.

### Behavioral Testing

#### Hindlimb Clasping Assay

To measure the hindlimb clasping phenotype previously described in(*42*), mice were suspended by their tail for a total of one min per trial. Four trials were conducted per mouse with 10 min between these trials. Scores were given based on the presence or absence of hindlimb clasping phenotype and ranged from 0-3 with 0 meaning hindlimb clasping was not present, 1 meaning that one hindlimb came inward toward the midline of the body, 2 meaning that both hindlimbs were pulled into the midline of the body without touching, and 3 meaning that both hindlimbs were pulled into the midline of the body and clasped.

### Dystonia-Like Behaviors During Tail Suspension

To better understand the relationship between hindlimb clasping and dystonic behavior, in the first cohort of mice we also scored dystonia-like behaviors which included limb grabbing (grabbing one or both hindlimbs and holding on), forelimb clenching (tight forelimb clenching into a fist), and twisting (twisting the trunk of the body so that it was perpendicular to the feet), and we also scored time that the hindlimbs were together.

Trunk twisting was scored on a scale of 0 = no abnormal twisting, 1 = twisting present less than 30 seconds, 2 = twisting present more than 30 seconds but less than one min, 3 = twisting present for whole min. Limb grabbing was scored as grabbing onto one or both of the back limbs and holding on with the scale ranging from 0 = not present, 1 = less than 30 s, 2 = more than 30 seconds, 3 = one full min. Finally, hindlimb clasping time was rated on the following scale: 0 = not present, 1 = only one limb at a time crossing midline, 2 = two limbs crossed midline less than 30 seconds, 3 = two limbs crossed midline more than 30 seconds.

#### Ledge Test

In order to quantify balance and coordination, the ledge test was performed in mice(*42*). Mice were placed onto the ledge of a cage so that their front and hind limbs were perched on the edge of the ledge. This task is scored from 0-3 with 0 meaning that there was no abnormal behavior and mice walked competently along the ledge without slips or walked normally and then lowered themselves into the cage competently, 1 meaning that mice slipped a few times with one hind limb but still managed to walk competently along the ledge, 2 meaning that mice dragged themselves forward across the ledge by their front paws with the hind paws below the ledge, and 3 meaning that the mice failed to move forward, lost their balance and nearly fell, or fell off of the ledge completely.

#### Erasmus Ladder

The Erasmus ladder is a commercially available apparatus from Noldus Technologies, which consists of higher and lower rungs was used to quantify gait deficits in this experiment. Animals ran on the Erasmus ladder for a total of 42 trials on each test day for 4 test days. The intertrial interval is varied between 10-20 s on each trial. A trial consists of a light turning on for 5 s and then wind from an air compressor turning on to entice the mouse to run. The mouse exits the chamber and runs to the chamber on the other side. Because there was no difference between days of Erasmus ladder runs, runs were averaged for all trials for each mouse.

### Ex vivo electrophysiology

Mice underwent a brief behavioral battery consisting of ledge test, hindlimb clasping , and cylinder test. Following testing, mice were anesthetized with isoflurane and were transcardially perfused with an ice-cold N-Methyl-D-Gluconate solution (osmolarity between 300-310 milliosmoles), which was bubbled in carbogen (5% carbon dioxide in 95 % oxygen). Brains were extracted and sliced sagitally into 300 um slices on a Leica vibratome at a rate of 0.1 – 0.5 μm/s. Slices were separated by hemisphere and transferred to a solution of NMDG warmed to 33 degrees C and then were incubated for 10-12 min. Slices were then transferred to a room temperature aCSF + sodium ascorbate solution, where they remained for at least one hour prior to recording. Slices were placed in a bath constantly infused with oxygenated aCSF for recording. K gluconate internal solution was vortexed for 30 min with amphoteriicng B and spun down for one min on a bench top centrifuge before being placed on ice. Microfill tips with Nalgene filters were attached to a 1 mL syringe filled with the amphotericin B infused K gluconate internal. Microfill tips were used to prefill fire-polished glass pipets with resistance 5-7 mOhms. A slice hold-down was placed and the dorsolateral striatum was located. We then performed cell-attached perforated patch on spiny projection neurons. Once successful perforation had occurred, we switched to current clamp and applied current steps from -350 to 600 by increments of 50, each interval occurred for 1 s every 10 s. We measured resting membrane potential and rheobase current (the amount of current required to elicit an action potential) for each cell.

### Immunohistochemistry

After transcardial perfusion with ice-cold 0.1 M Phosphate buffered saline followed by 4% paraformaldehyde in 0.1 M Phosphate buffered saline, brains were extracted and placed overnight in PFA. The next day, brains were transferred to a 30% sucrose solution until they sunk. Brains were then cut on the vibratome into 50 μm sections. As previously done (*11*)slices were washed in 0.1 M PBS between each step. Slices were first blocked with 0.3% horse block and 1% triton-x100 in PBS (hereafter blocking solution). Slice were then incubated overnight at 4⁰C in primary antibody cocktail for *Gnal,* cyclic adenosine monophosphate, and a primary antibody for phosphorylated protein kinase A. Thereafter, slices were put in a secondary antibody cocktail for 2h at room temperature and incubated with DAPI for 10 min before being mounted onto histobond slides for further analysis.

### Data Analysis

All data were analyzed in GraphPad Prism 10.0.1 with alpha set to 0.05. Two-tailed unpaired t-tests were used to evaluate differences between two groups. For analysis of hindlimb clasping and ledge test a non-parametric t-test (Mann-Whitney U) was used. For analyses comparing 3 or more groups, a one-way ANOVA was used, or in cases when hindlimb clasping or ledge test data were evaluated, a non-parametric Kruskal-Wallis test was used. In cases where within-subjects hindlimb clasping data were compared, non-parametric Friedman ANOVA was used. When appropriate to evaluate differences between groups, Tukey post-hocs were used for parametric ANOVAs, and Dunn’s post-hocs were used for non-parametric ANOVAs. Electrophysiology data were first analyzed in pClamp 11 software. Subsequent analyses were conducted in GraphPad Prism. We used independent sample t-tests to evaluate differences between two groups, and we used a mixed-effects model to analyze differences in response to current steps. For correlations between hindlimb clasping and dystonia-like behavior, non-parametric Spearman’s R tests were used.

## Funding

National Institutes of Health grant R00NS110878 (MM)

Tyler’s Hope for a Dystonia Cure (MM)

## Author contributions

Conceptualization: NC, MM

Methodology: NC, DH, SB, EM, TC, MK, SG, IG, DN, KH, MM, MM

Investigation: NC, SB, TC, MK, SG, IG

Funding acquisition: MM

Project administration: NC, MM

Supervision: NC, MM

Writing – original draft: NC, MM

Writing – review & editing: NC, DH, SB, EM, TC, MK, SG, IG, DN, KH, MM, MM

## Data and materials availability

All data, analysis, and analysis code are available from the corresponding author upon request. Unique materials generated as part of this manuscript are subject to an MTA.

