## Supplemental Figures for "Conditional Knockout of Striatal Gnal Produces Dystonia-like Motor Phenotypes"

**Supplementary Materials**

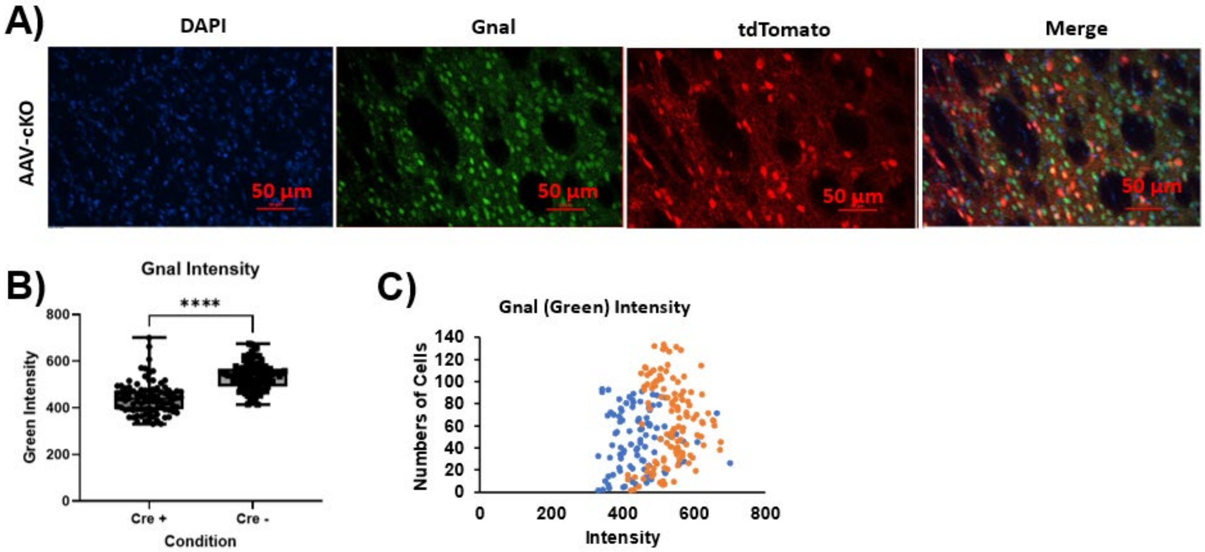


**Figure S1. AAV-cKO virus leads to knockout of *Gnal*.** A) From left to right, max intensity images from the striatum of an AAV-cKO homozygous mouse, in blue, DAPI, in green, Gnal, in red, tdTomato, and a merged image showing these images overlayed. B) Shows the overall fluorescent intensity of green (Gnal) in cells positive for Cre (Cre+) and cells negative for Cre (Cre-). Overall, a two-tailed independent samples t-test ($\alpha = 0.05$) showed that cre + cells (n = 94) had significantly less green fluorescent intensity than Cre – cells (n = 121), demonstrating loss of Gnal in cells which were transduced with the AAV-cKO virus. C) Shows the overall fluorescent intensity of green (Gnal) in the Cre + and Cre – populations of cells overlaid to compare the two conditions. Dots representing fluorescent intensity are displayed in blue for Cre + cells and in orange for Cre – cells.

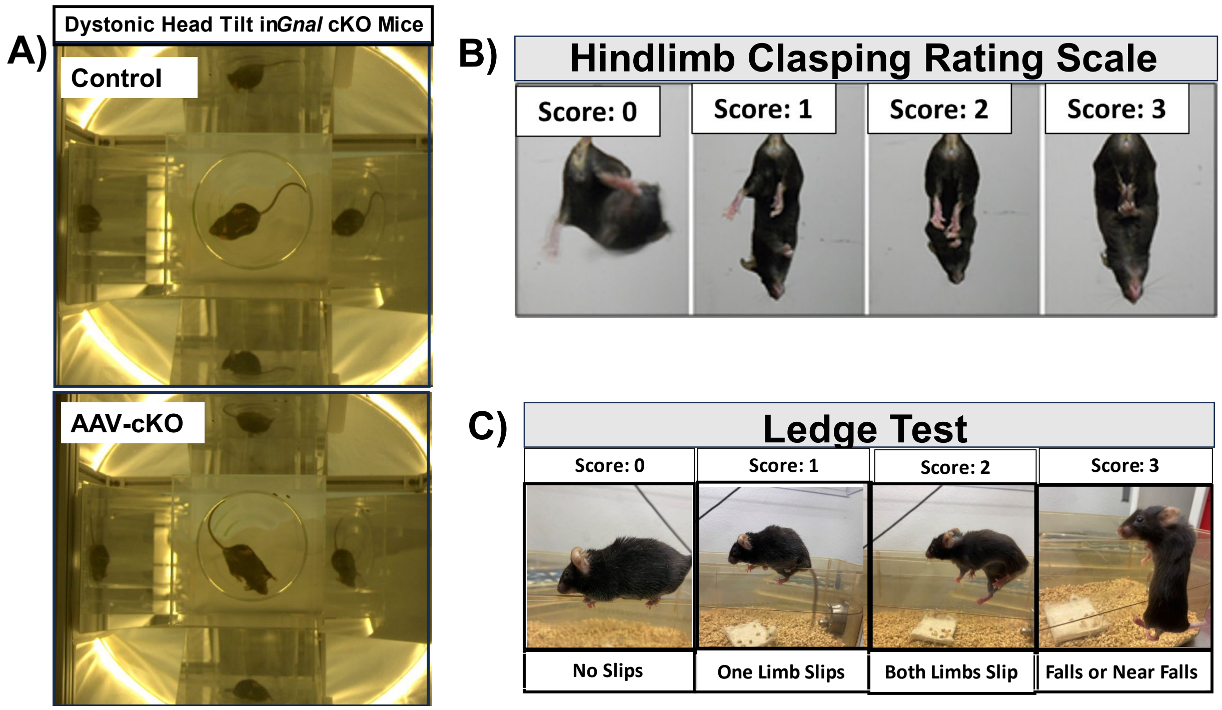


**Figure S2. Rating Scales and Dystonia-like Head Tilt Observed in *Gnal* cKO Mouse.** A) 3-D Neurobehavioral Chamber (3DNC) which was designed and built by Nicole Chambers for use in mouse experiments. Images show the control mouse (above) with no head tilt, and the *Gnal* cKO mouse (below) with significant head tilt. In human *GNAL*-linked dystonia, torticollis is a main symptom. B) Rating scale used for hindlimb clasping, rated as follows: 0 = no hindlimb clasping present; 1 = one foot brought inward and crossing the midline; 2 = two hindlimbs were brought inward toward the midline but not touching; 3 = two hindlimbs were brough inward toward the midline and clasped. C) Rating scale used for the ledge test, which is scored as follows: 0 = mouse walked competently along the ledge with no slips and/or lowered itself competently into the cage, 1 = one foot at a time slipped off the ledge but the mouse otherwise walked competently, 2 = both of the mouse's hindlimbs slipped down from the ledge and the mouse pulled itself forward with the front limbs; 3 = mouse slipped and fell or nearly fell off of the ledge.

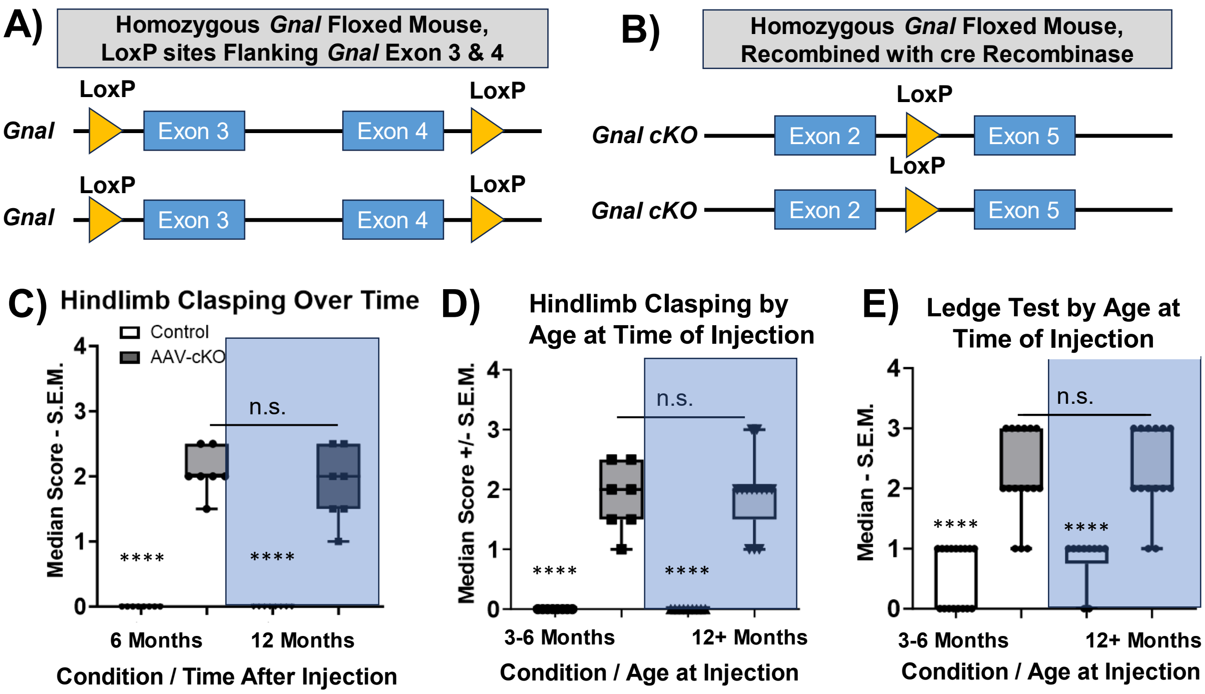


**Figure S3. Dystonia-Related Deficits are Present in Homozygous AAV-cKO Mice Regardless of Age** A) Shows the genetic mutation present in our *Gnal* floxed animals. In this case, in a homozygous *Gnal*-floxed animal, both copies of *Gnal* contain a LoxP site flanking exons 3 and 4 of the gene. B) Shows the consequences of recombination of cre-recombinase. One LoxP site remains and genetic information between exon2 and exon5 is deleted. This results in a nonsense deletion of the *Gnal* gene (AAV-cKO). Data in S3C-E are analyzed via Kruskal-Wallis ANOVA ($\alpha = 0.05$), each dot represents one mouse. C) Data from the same mice at 6 months and 12 months after injection. Overall, once dystonia-like symptoms develop, they remain stable over time. D) Shows data from two cohorts of *Gnal* knockout mice, showing that regardless of the age at time of injection, mice develop stable dystonia-like symptoms of a similar magnitude. E) Shows data from two cohorts of *Gnal* knockout mice, either 3-6 months or 12+ months of age at the time of injecgtion. Overall, there is no difference in ledge test deficit present in AAV-cKO mice regardless of age at time of injection.

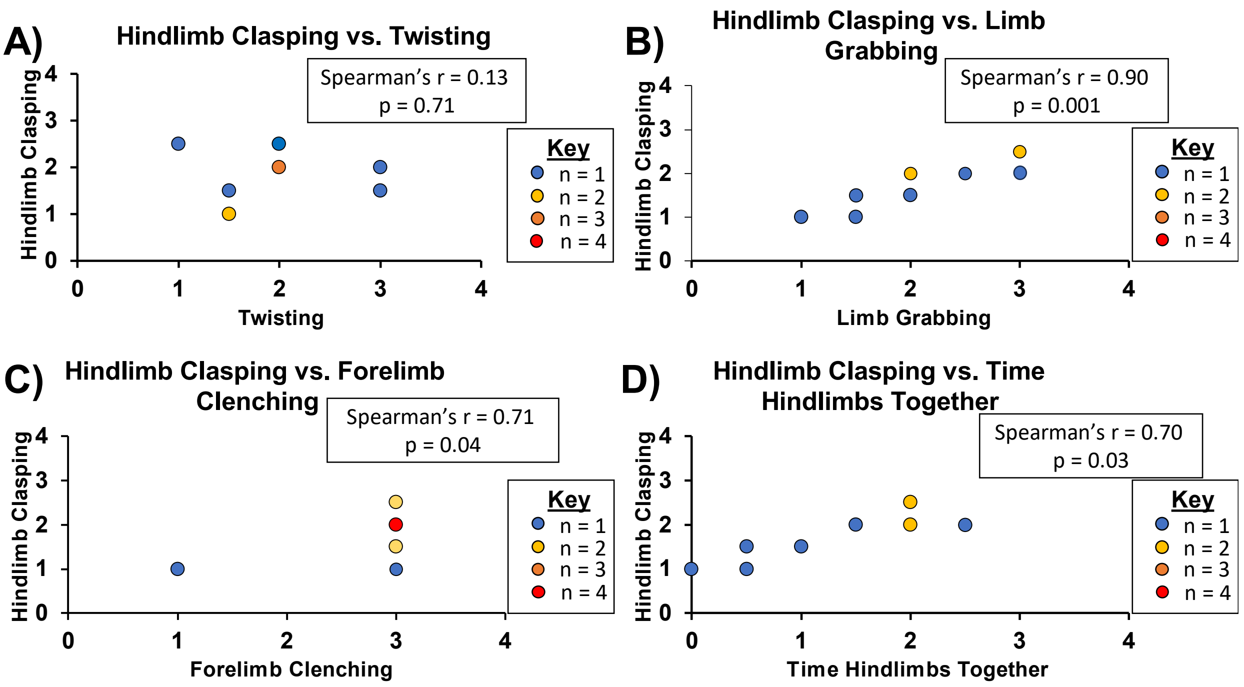


**Figure S4. Hindlimb Clasping Behavior and Dystonia Behaviors Are Correlated During Tail Suspension.** A) Shows medians for AAV-cKO animals for hindlimb clasping and twisting behaviors during baseline. Overall, there is no correlation between hindlimb clasping and twisting. B) Shows medians for AAV-cKO animals for hindlimb clasping and limb grabbing behavior. Overall there is a strong positive correlation between these two variables. C) Shows median scatterdot of AAV-cKO animals for hindlimb clasping and forelimb clenching behavior. Overall, there is a significant positive correlation between hindlimb clasping and forelimb clenching. D) Shows median scatterdot of AAV-cKO animals for hindlimb clasping and time hindlimbs together ( 0 = not together, 1 = just one limb in toward midline <30s, 2 = Both hindlimbs in toward midline < 30s, 3 = both hindlimbs toward midline > 30s). N = 10 for all, not all dots appear on the graphs due to overlap in scores. In these cases, the dot is color-coded yellow if 2 animals, orange if 3 animals, red if 4 animals. All data were analyzed using Spearman's non-parametric correlation, α = 0.05.

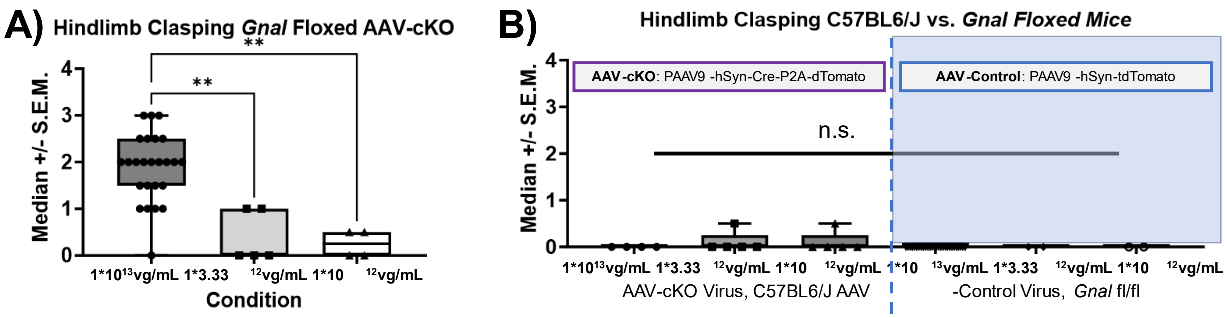


**Figure S5. AAV-cKO virus does not cause off-target effects in C57BL6/J mice.** In each case data were analyzed via Kruskal-Wallis ANOVA with Dunn post-hoc tests, $\alpha = 0.05$. Each dot represents one mouse. A) Medians of hindlimb clasping scores are displayed for each viral dilution injected into homozygous *Gnal* floxed animals. Overall, only the $1*10^{13}$ vector genomes per milliliter (vg/mL) concentration produced hindlimb clasping in *Gnal* floxed animals. B) Medians of hindlimb clasping for C57BL6/J mice receiving AAV-cKO virus on the left and *Gnal* floxed mice receiving control virus on the right. Overall, there were no differences in hindlimb clasping between C57BL6/J mice receiving AAV-cKO virus and *Gnal* floxed mice receiving the control virus, suggesting that the effects that we observed in this paper are due to *Gnal* knockout and not due to random, off-target effects.

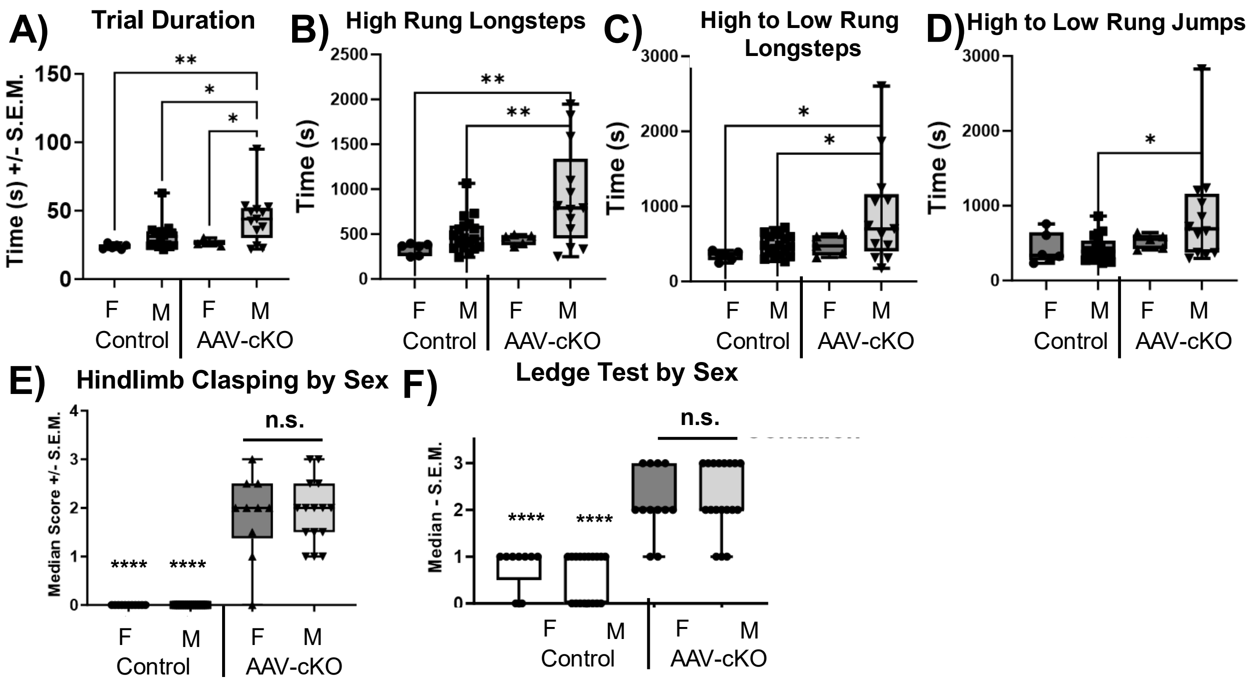


**Figure S6. Sex Differences Observed in Erasmus Ladder Task and Exploring Effects of Viral Dilution on Phenotype.** All data in this figure were analyzed via One-Way ANOVA, α = 0.05. Each dot represents averaged data for 42 trials for one mouse. A) Shows the average of 42 trials +/- S.E.M. for trial duration in the Erasmus ladder, separated by sex and condition. F is Female, M is male. Overall, female and male controls run faster than AAV-cKO males, but not than AAV-cKO females. B) Average time to complete high rung longsteps, averaged for each mouse over female and male controls took less time to complete high rung longsteps than male AAV-cKO animals. There is no difference between controls and female AAV-cKO mice.C) Average time to complete high to low rung longsteps. Female and male controls less time to complete H-L steps than male AAV-cKO mice but not female AAV-ckO mice. D) Shows time to complete High to low rung jumps. Male control mice took less time to complete jumps than male AAV-cKO mice. There were no differences between female control and female AAV-cKO mice. Overall these data suggest a beneficial effect of female sex on functional deficits of dystonia.

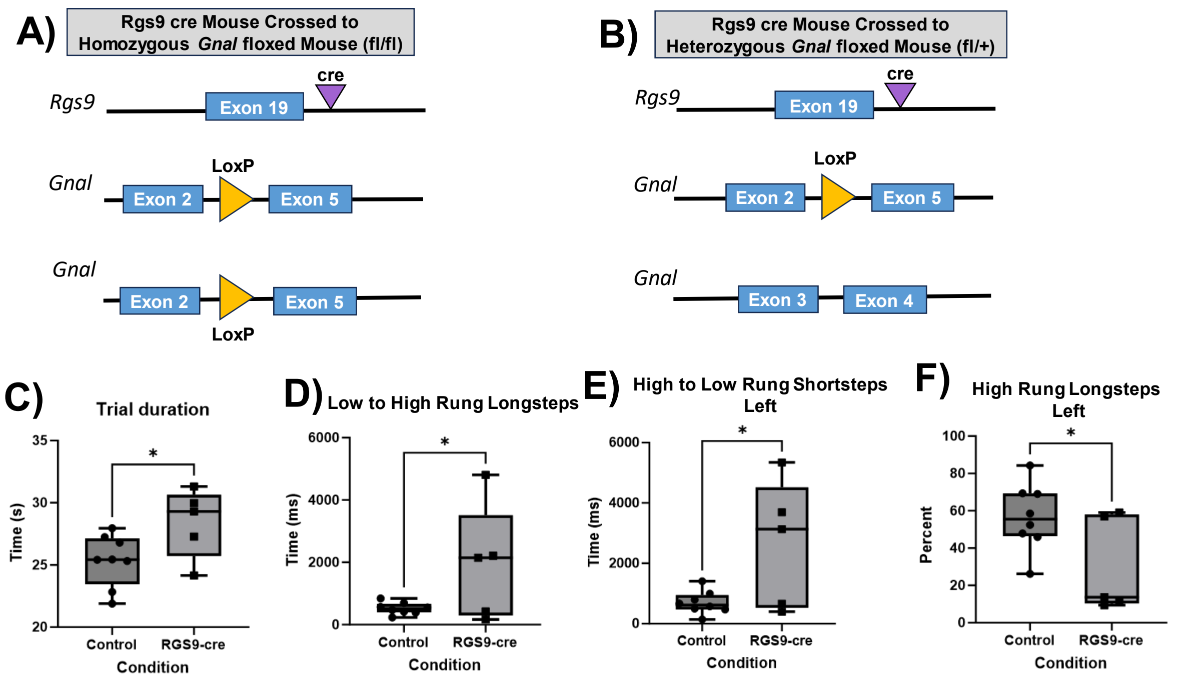


**Figure S7. RGS9-cre Mice Show Similar Deficits to AAV-cKO in Erasmus Ladder.** A) Shows the consequences of knocking out both copies of *Gnal* in homozygous *Gnal* floxed mice. B) Shows the consequences of cre-recombination in heterozygous *Gnal* floxed mice, where only one copy of *Gnal* is likely knocked out. All data from Erasmus ladder were tested using independent samples t-test with alpha set to 0.05. Data represent time to complete different tasks averaged over 42 trials. Data are collapsed across RGS9 cre conditions (n = 2 homozygous Gnal floxed RGS9-cre, 3 heterozygous Gnal floxed RGS9-cre C) Shows that trial duration is increased in RGS9-cre animals. D) Shows that time to complete low to high rung longsteps is increased in RGS9-cre mice. E) Shows that percentage of high rung steps is decreased in RGS9-cre animals. F) Shows that the time to complete High to low rung shortsteps on the left is increased in RGS9 cre mice. Overall, these deficits mirror those seen in AAV-cKO animals.

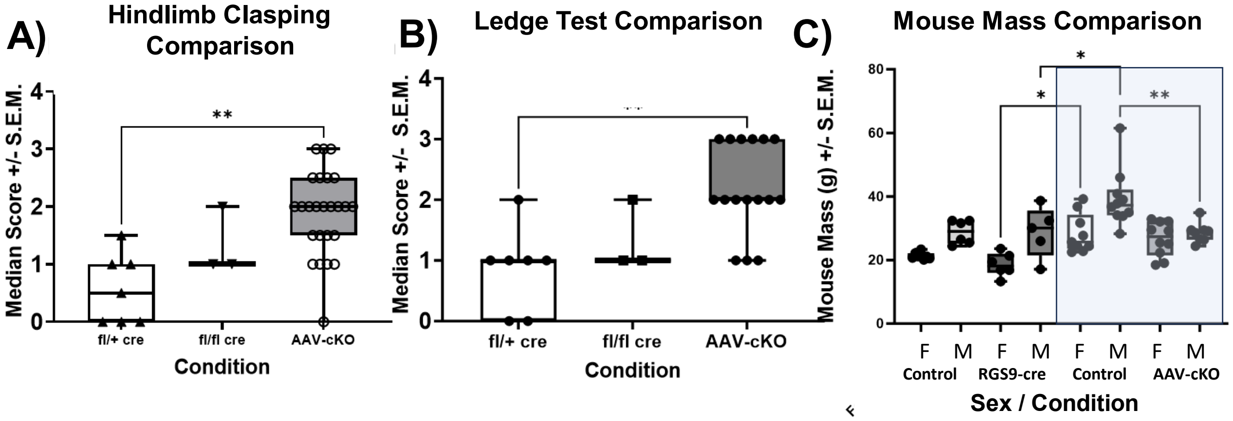


**Figure S8. Comparison of Viral Striatal cKO mice and embryonic *Gnal* Knockout.** A) Hindlimb clasping scores were compared for heterozygous (fl/+ cre) and homozygous (fl/fl cre) mice positive for *Rgs9* Cre, and viral striatal *Gnal* cKO mice (AAV-cKO). Data were analyzed using Kruskal-Wallis test, α = 0.05. *Gnal* fl/+ heterozygous mice show less hindlimb clasping than AAV-cKO mice. Homozygous mice positive for *Rgs9* cre and AAV-cKO mice are similar in hindlimb clasping. B) Ledge test comparisons revealed that heterozygous Rgs9 Cre mice show more coordination on ledge test than AAV-cKO mice, whereas homozygous Rgs9 Cre mice are similar to AAV-cKO. C) Mass comparison between heterozygous and homozygous Rgs9 Cre mice and homozygous AAV-cKO mice. Data for homozygous and heterozygous mice are collapsed for Rgs9 Cre mice. Rgs9 Cre positive females and males weighed less than sex matched AAV controls, suggesting that the Rgs9 Cre mice are feeding less or that they have metabolic issues. Another interesting finding from these data was that male AAV-cKO animals weigh less than male controls, likely due to the dystonic symptoms. Only significance points representing comparisons across the same sex were retained for this graph. * $p < 0.05$, ** $p < 0.01$

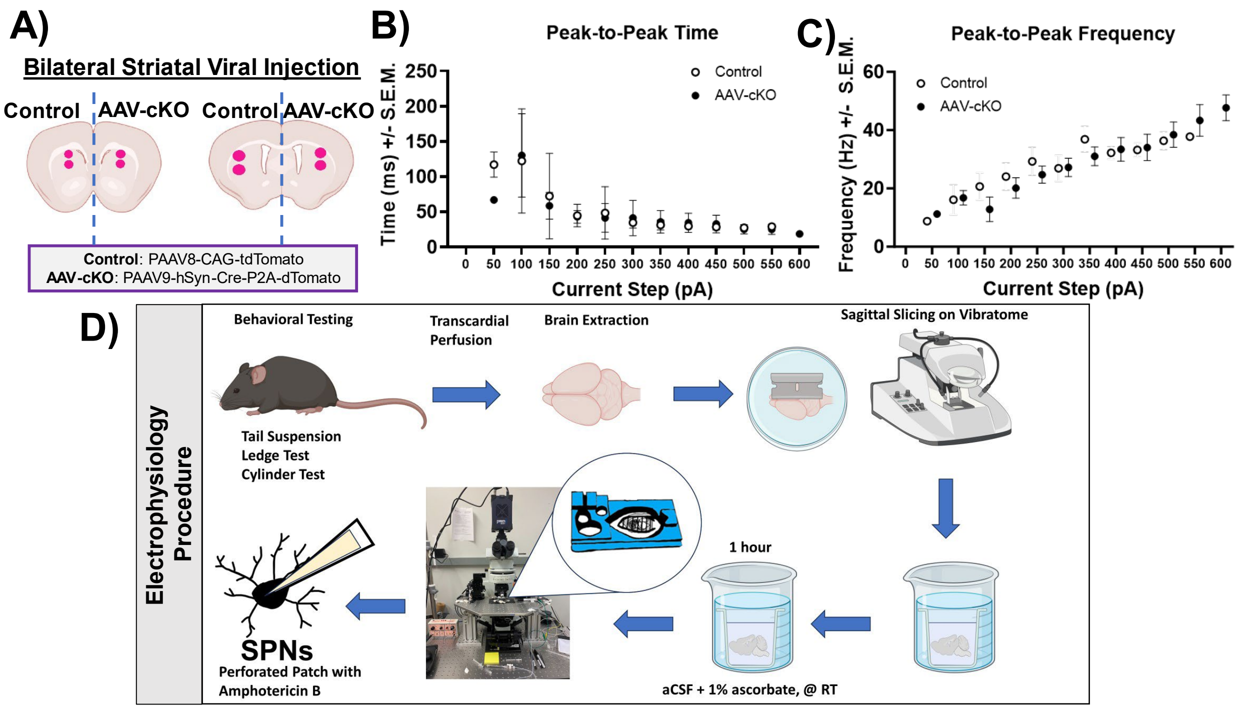


**Figure S9. Peak-to-peak time or frequency were similar in control and AAV-cKO striatal spiny projection neurons.** A) For electrophysiology experiments, mice were injected with the control virus in the left striatum and the AAV-cKO virus in the right striatum. B) There were no differences between control and AAV-cKO cells in peak-to-peak time, Data in S9B and S9C were analyzed via mixed effects model, α = 0.05. C) There were no differences in peak-to-peak frequency between control and AAV-cKO cells. D) Shows the procedure used on electrophysiology test days. Animals underwent a brief behavioral battery. Animals were anesthetized with isoflurane and transcardially perfused with N-methyl-D-gluconate (NMDG). The brain was then extracted and cut into sections using a vibratome in oxygenated NMDG. Slices were moved to oxygenated NMDG at 34 degrees C for 10-12 min before being moved to an oxygenated artificial cerebrospinal fluid (aCSF) containing 1% sodium ascorbate. Slices were incubated for 1 hour at room temperature before being transferred to the bath chamber which was constantly perfused with oxygenated aCSF. Cells expressing the virus were identified according to the expression of fluorescent marker and then recorded with a sharp glass pipette containing Amphotericin B in K-Gluconate internal solution.

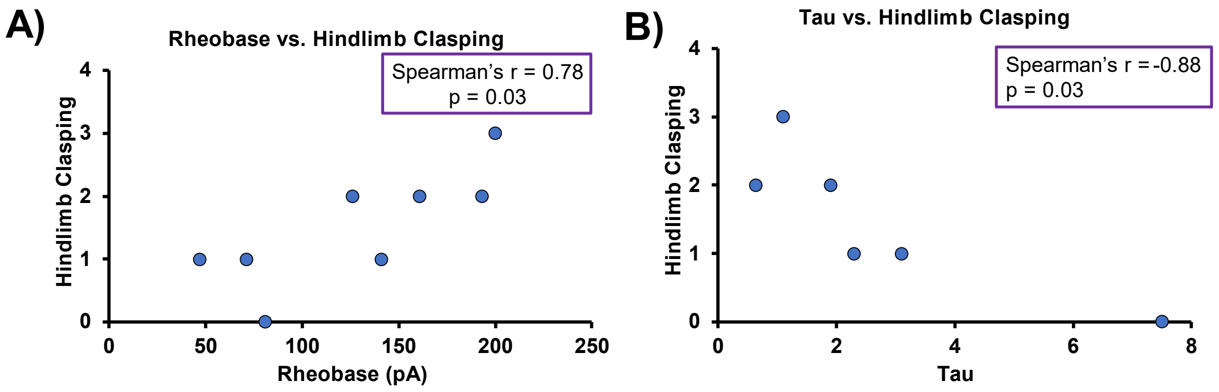


**S10. Altered Electrophysiological Properties are Correlated with Hindlimb Clasping Dystonic Phenotype.** Data in A and B were correlated using non-parametric Spearman correlation, α = 0.05. A) Data from hindlimb clasping in AAV-cKO mice were correlated for those mice that underwent behavior and had cells that were tested for rheobase during electrophysiology (n = 8). Overall, there was a positive correlation where rheobase weas increased in mice showing more hindlimb clasping. B) Data from hindlimb clasping in AAV-cKO mice was correlated with tau in AAV-cKO mice that underwent behavior and had measurements for tau (n = 6). Overall, there was a negative correlation for tau where the more hindlimb clasping the smaller the tau (ms) and the less hindlimb clasping the larger the tau.
